# Avatar-Induced Postural Responses during a Virtual Full-Body Illusion

**DOI:** 10.64898/2026.09.09.750263

**Authors:** Yuki Tsuji, Kenichiro Furuya, Katsuki Higo, Sotaro Shimada

**Affiliations:** Department of Complex and Intelligent Systems, School of Systems Information Science, Future University Hakodate, Hakodate, Japan; Department of Electrical Engineering, Graduate School of Science and Technology, Meiji University, Kawasaki, Japan; Organization for the Strategic Coordination of Research and Intellectual Properties, Meiji University, Kawasaki, Japan; Department of Electronics and Bioinformatics, School of Science and Technology, Meiji University, Kawasaki, Japan

**Keywords:** **Keywords**: Full-body illusion, Virtual reality, Body ownership, Postural control, Center of pressure

## Abstract

Body ownership over an artificial body can alter perceptual and autonomic responses, but its influence on online postural control remains less understood. We investigated whether ownership-related body–avatar correspondence modulates postural responses to an unexpected change in avatar posture. Thirty-six healthy male participants were randomly assigned to an Embodiment condition, in which body–avatar correspondence was maintained, or a Disruption condition, in which the avatar’s arm temporarily moved independently of the participant before the critical event. During a standing virtual-reality session with synchronous visuotactile stimulation, the avatar unexpectedly leaned forward. We introduced Avatar-Induced Postural Response (AIPR), an event-related index of anterior–posterior center-of-pressure displacement following the avatar’s movement. Ownership-related questionnaire ratings were higher in the Embodiment condition, whereas self-location-related ratings did not differ between conditions. AIPR was greater in the Embodiment condition and was greater than zero only in this condition. AIPR was also positively correlated with skin conductance responses in the Embodiment condition, but was not associated with questionnaire ratings. These findings suggest that an unexpected change in avatar posture can influence the participant’s own postural regulation when ownership-related body–avatar correspondence is maintained. AIPR may provide a complementary behavioral index of a postural-control component of avatar embodiment not fully reflected in explicit subjective reports.

## Introduction

Body representation underlies the perception and control of one’s own body. It is constructed from multiple sources of body-related sensory information, including vision, touch, and proprioception. These representations are not unitary: some contribute to the conscious perception and recognition of the body, whereas others support the online control of posture and action (Dijkerman & de Haan, 2007; de Vignemont, 2010). By integrating sensory information within body-centered reference frames, body representations enable individuals to locate their body and its parts in space and to coordinate bodily actions in relation to the surrounding environment (Blanke et al., 2015). They thus provide the basis for perceiving and controlling the body during interactions with the external world.

Body representations are not static but can be modified in response to multisensory information. Experimental manipulations of the correspondence between visual, tactile, and proprioceptive signals can alter the experience of body ownership, demonstrating the plasticity of body representation (Kilteni et al., 2012). Immersive virtual reality (VR) makes it possible to systematically manipulate the visual perspective, appearance, and multisensory contingencies of a virtual body, thereby enabling controlled investigations of the conditions under which body ownership emerges (Slater et al., 2010; Maselli & Slater, 2013). VR therefore provides a powerful experimental framework for manipulating body representation and investigating the multisensory mechanisms underlying body ownership.

The rubber hand illusion (RHI) is one of the most widely used paradigms for investigating the plasticity of body ownership. In the classic RHI, synchronous stroking of a visible rubber hand and the participant’s hidden real hand can induce the experience that the rubber hand belongs to one’s own body (Botvinick & Cohen, 1998). The RHI has typically been assessed using subjective questionnaires together with proprioceptive drift, defined as a shift in the perceived location of the real hand toward the rubber hand. Physiological responses have also been examined, including skin conductance responses to threats directed toward the rubber hand and changes in the skin temperature of the real hand (Armel & Ramachandran, 2003; Moseley et al., 2008). However, these measures do not necessarily reflect a unitary process. Subjective ownership and proprioceptive drift may be weakly associated but can exhibit different sensitivities and response patterns to visuotactile incongruence, suggesting that they reflect at least partly distinct processes (Rohde et al., 2011; Shimada et al., 2014). The RHI has therefore established a framework in which body ownership is examined using complementary subjective, behavioral, and physiological measures.

Research on body ownership has subsequently been extended from individual body parts to the whole body through the full-body illusion (FBI). In a typical FBI paradigm, participants view a real or virtual body from a first- or third-person perspective while receiving synchronous visuotactile stimulation, which can induce the experience that the observed body belongs to them and alter the perceived location of the self (Lenggenhager et al., 2007; Ehrsson, 2007; Petkova et al., 2011). Full-body illusions have therefore been used to investigate multiple components of bodily self-consciousness, particularly body ownership and self-location. These components are commonly assessed using subjective questionnaires, behavioral estimates of self-location, and autonomic responses to events directed toward the observed body, such as skin conductance and heart-rate responses (Petkova & Ehrsson, 2008; Maselli & Slater, 2013). Importantly, body ownership and self-location do not necessarily vary together, indicating that they reflect partly dissociable aspects of bodily self-consciousness (Maselli & Slater, 2014). However, although these measures characterize perceptual, spatial, and autonomic consequences of full-body embodiment, they provide only limited insight into how embodiment influences the participant’s own motor regulation. In particular, relatively little is known about how ownership of an artificial whole body influences online postural control.

One potential consequence of embodiment is that changes occurring in an embodied artificial body may influence the participant’s own motor system. Shibuya et al. (2018) examined this possibility using a rubber hand illusion paradigm in which a participant’s hidden hand and a model hand displayed on a monitor were stroked either synchronously or asynchronously. When the stroking was unexpectedly interrupted by a movement of the model hand, participants more frequently exhibited spontaneous finger movements corresponding to the observed movement in the synchronous condition than in the asynchronous condition, despite being instructed to remain relaxed. Shibuya et al. interpreted this behavioral effect as motor back projection, whereby a movement attributed to an embodied artificial body is reflected back onto the participant’s corresponding real body part. This finding raises the possibility that a comparable process may extend beyond an individual limb to whole-body postural control in the full-body illusion. In particular, it remains unclear whether a change in an avatar’s posture induces a corresponding postural adjustment in a standing participant and whether this response is modulated by ownership-related body–avatar correspondence.

To examine this possibility, participants stood on a force plate while viewing an avatar from behind and receiving synchronous visuo-tactile stimulation. In the Embodiment condition, body–avatar correspondence was maintained until the critical postural event. In the Disruption condition, this correspondence was transiently violated beforehand when the avatar’s arm moved independently of the participant. Subsequently, the avatar unexpectedly leaned forward in both conditions without a corresponding movement initiated by the participant. If motor back projection extends to whole-body postural control, the avatar’s forward-leaning movement should elicit a corresponding anterior displacement of the participant’s COP, particularly when body–avatar correspondence has been maintained. To quantify this postural response, we introduced Avatar-Induced Postural Response (AIPR), a COP-based measure of the direction and magnitude of anterior–posterior postural displacement following the avatar’s movement relative to the pre-event baseline.

To determine how AIPR relates to established subjective and physiological manifestations of full-body embodiment, we also collected questionnaire ratings, SCRs, and heart-rate-related measures. Ownership- and self-location-related questionnaire ratings were used to determine whether the Disruption manipulation affected the subjective experience of ownership-related body–avatar correspondence and whether this effect extended to perceived self-location, which can be partly dissociated from body ownership (Lenggenhager et al., 2007; Maselli & Slater, 2014). SCRs were recorded to assess event-related sympathetic responses to the avatar’s movements, because events involving an embodied artificial body can elicit changes in electrodermal activity (Petkova & Ehrsson, 2008; Maselli & Slater, 2013). Instantaneous heart rate and the LF/HF ratio were examined as exploratory indices of cardiovascular responses to the unexpected avatar movements, given that salient or unexpected events can evoke rapid changes in heart rate and autonomic activity (Slater et al., 2010; Bradley et al., 2012; Noordewier et al., 2021). We further examined the associations of AIPR with questionnaire ratings and SCRs to determine whether the postural response covaried with explicit subjective experience or autonomic reactivity, or instead captured a partly distinct aspect of avatar-related embodiment. We hypothesized that AIPR values would be greater in the Embodiment condition than in the Disruption condition.

## Materials and Methods

### Participants

A total of 36 healthy male participants took part in the study and were included in the analysis (mean age = 21.5 years, SD = 1.1). An additional three participants did not complete the experiment because of cybersickness and were therefore excluded. All participants were students at Meiji University or young adults living near the university. Because the present study used a male avatar and focused on an initial validation of AIPR, only male participants were recruited to keep avatar–participant gender correspondence constant. The study procedures were approved by the ethics committee of Meiji University (24-574), and the study was conducted in accordance with the principles and guidelines of the Declaration of Helsinki. All participants provided written informed consent and received compensation for their participation.

### Apparatus

Participants wore a head-mounted display (HMD; HTC Vive, HTC, Taiwan) and two motion trackers (HTC Vive Tracker, HTC, Taiwan) attached to the left and right wrists. They were instructed to remove their shoes and stand comfortably on a portable force plate (TFG-4060, Tech Gihan Co., Ltd., Japan), with their arms hanging naturally alongside the body. The force plate was used to record center of pressure (COP). Participants were also instructed to remain as relaxed as possible during the experiment.

The virtual environment was created using Unity 2019.2.2f1. Participants viewed the back of an avatar through the HMD. The avatar’s movements were controlled using three-point tracking based on the HMD and the two wrist trackers. Before the experiment, the height and shoulder width of the avatar were adjusted based on each participant’s self-report so that the avatar approximately matched the participant’s body dimensions. The location on the avatar’s back corresponding to tactile stimulation was also adjusted to match the participant’s perceived back position.

A VR controller (HTC Vive Controller, HTC, Taiwan) was used to synchronize tactile stimulation with visual stimulation in the virtual environment. A stick used to stroke the participant’s back was fixed to the VR controller. This setup allowed the experimenter’s stroking movements on the participant’s back to be presented as corresponding movements of a virtual stick touching the avatar’s back. The experimental setup and VR view are shown in Fig. 1A.

**Fig. 1.**
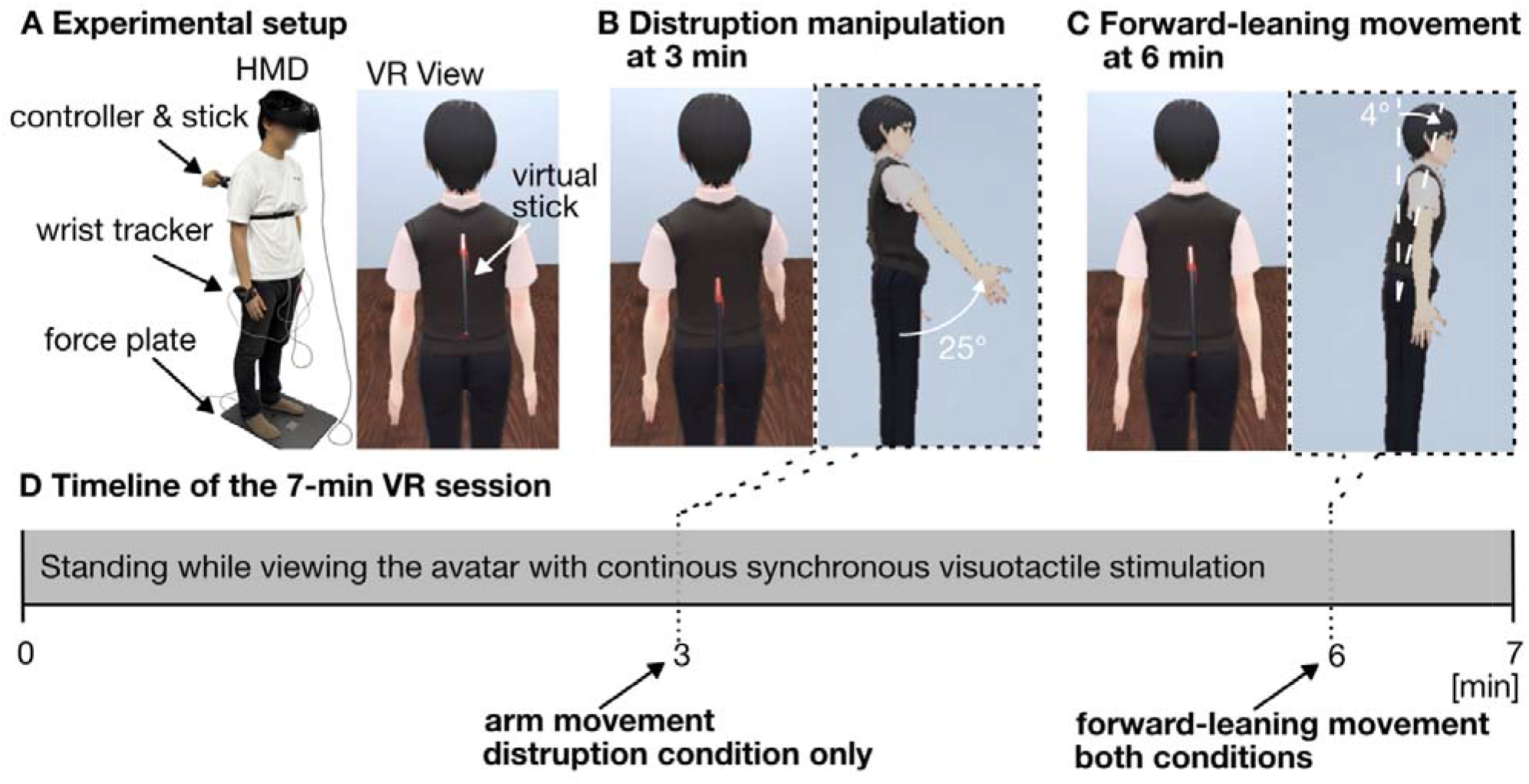
**Experimental setup, avatar movements, and experimental timeline. A**: Participants stood on a force plate while wearing an HMD and wrist trackers. The experimenter used a controller-mounted rod to stroke the participant’s back, while the VR view showed a virtual stick touching the back of the avatar. An author posed as the participant for the photograph in panel A and consented to its publication. **B**: In the Disruption condition, the avatar’s right arm was temporarily decoupled from the participant’s wrist tracking at 3 min and moved forward independently. **C**: In both conditions, the avatar was temporarily decoupled from the participant’s tracking at 6 min and performed a forward-leaning movement. **D**: Timeline of the 7-min VR session. After the session, participants completed the questionnaire.

Electrocardiogram (ECG) and electrodermal activity (EDA) were recorded continuously during the experimental session using a biosignal acquisition system (BITalino (r)evolution, PLUX Wireless Biosignals S.A., Portugal). ECG, EDA, and COP data were sampled at 1000 Hz, with COP data recorded simultaneously using the force plate.

For ECG and EDA measurements, wired biosensors were used, and electrodes were directly affixed to the skin using self-adhesive Ag/AgCl gel electrodes (Arbo™, Kendall™, Cardinal Health™, USA) to reduce contact resistance. ECG was recorded in a Lead II configuration, with electrodes placed at the left sixth intercostal space (IN+), below the left clavicle (REF), and below the right clavicle (IN-). For EDA, electrodes were placed on the proximal phalanges of the index (IN+) and middle (IN-) fingers of the right hand. All physiological sensors were connected to the BITalino system via a development kit, and data were transmitted to a PC via Bluetooth.

Experimental events were synchronized across the VR environment, physiological recordings, and COP recordings using trigger signals transmitted from the VR controller through an Arduino Uno R3.

### Experimental design and procedure

The study used a randomized between-subjects design to examine whether ownership-related body–avatar correspondence modulated participants’ postural responses to an unexpected change in avatar posture. Participants were assigned to either the Embodiment condition or the Disruption condition, with 18 participants in each condition. The primary behavioral outcome was AIPR, operationalized as the baseline-standardized event-related anterior–posterior COP displacement following the avatar’s forward-leaning movement. Questionnaire ratings and autonomic measures were included as complementary indices of avatar-related embodiment.

Before the experiment, participants were informed about the general procedure, the equipment, the experimental task, and their rights as participants. They were told that they could pause or terminate the session at any time if they experienced discomfort, including VR sickness. To minimize response bias, the specific hypotheses and the differences between the experimental conditions were not disclosed.

Participants completed a single continuous 7-min standing VR session. Throughout the session, they were instructed to remain relaxed, maintain their standing posture, and observe the avatar. To induce avatar-related embodiment in both conditions, the experimenter continuously stroked the participant’s back with the controller-mounted rod while the virtual stick was shown touching the corresponding location on the avatar’s back.

The two conditions differed only in the event presented 3 min after the start of the session. In the Disruption condition, the avatar’s right arm was temporarily decoupled from the participant’s wrist tracking and executed a predefined 6-s forward movement independently of the participant’s actual wrist movement. Starting from its resting position alongside the body, the arm rotated forward about the shoulder joint by approximately 25°, remained briefly in the forward position, and then returned to its original position (Fig. 1B). Wrist tracking resumed immediately after the movement. No corresponding arm movement occurred in the Embodiment condition.

In both conditions, 6 min after the start of the session, the avatar was temporarily decoupled from the participant’s tracking and performed an unexpected 3.5-s forward-leaning movement that was not initiated by the participant. The avatar leaned forward, remained briefly in the forward-leaning posture, and then returned to its original upright posture (Fig. 1C). This event served as the trigger for calculating the AIPR from the participant’s COP data.

After the 7-min VR session, participants completed the questionnaire assessing avatar-related bodily experiences. COP, ECG, and EDA were recorded continuously throughout the session. The sequence of experimental events is summarized in Fig. 1D

### Questionnaire

Questionnaire responses were collected after the experimental session. The questionnaire items were translated into Japanese based on the questionnaire used by Lenggenhager et al. (2007) and were partially modified to suit the present experimental context (Table 1). Participants responded to each item using a 7-point Likert scale ranging from strongly disagree (−3) to strongly agree (+3).

**Table 1.**
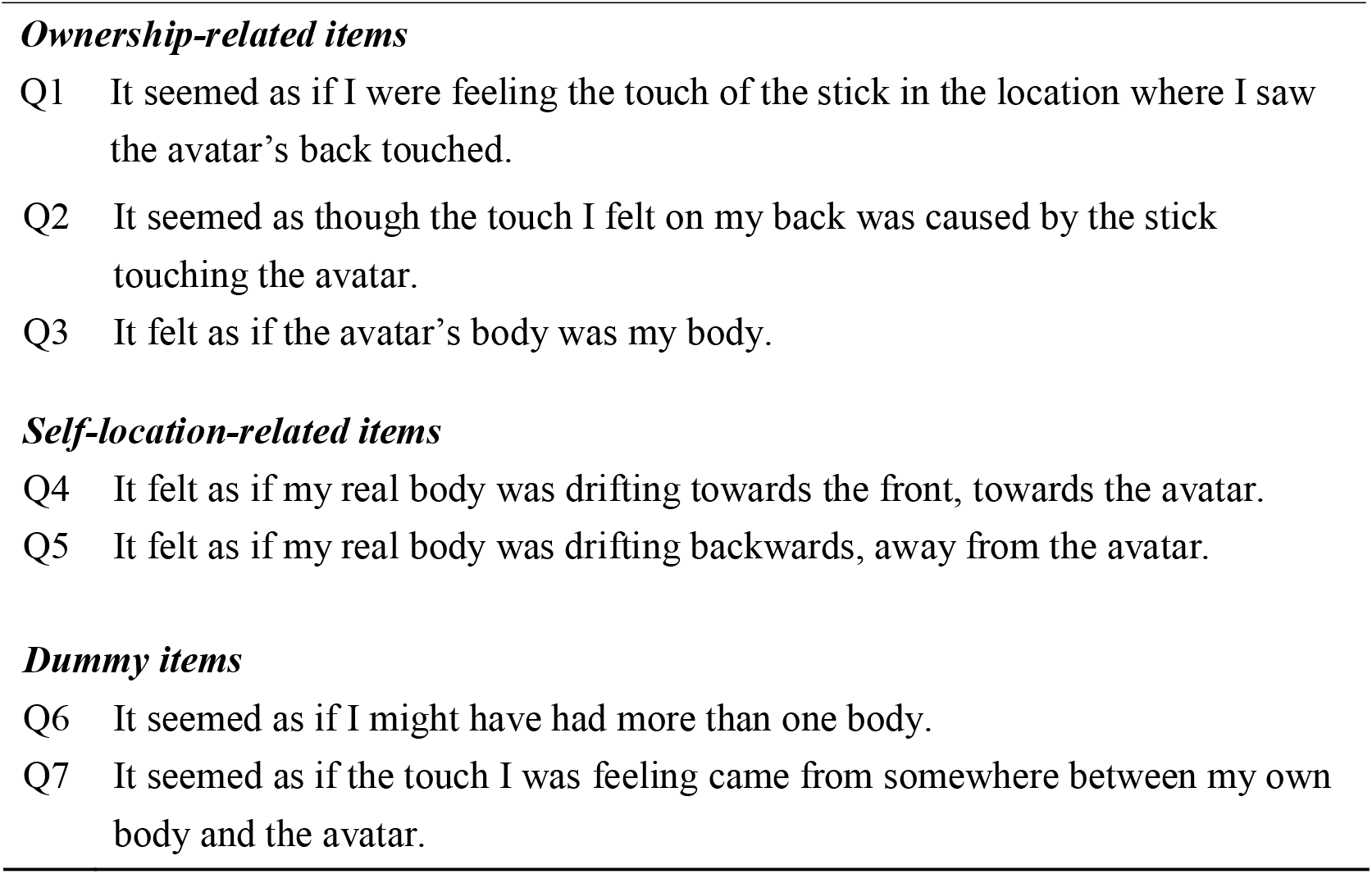
**Questionnaire items used in the present study**

The questionnaire administered immediately after the experimental session consisted of seven items. Items 1–3 assessed body ownership, items 4–5 assessed self-location, and items 6–7 served as dummy items.

### Data analysis

#### Center-of-Pressure (COP)

Center-of-pressure (COP) data were analyzed as an index of postural control because force-plate-derived COP trajectories are widely used to quantify postural control during standing (Quijoux et al., 2021). Because the present study focused on forward and backward postural responses to the avatar’s forward-leaning movement, the anterior–posterior component of the COP signal was used for analysis. The COP time series was first smoothed using a moving average with a window size of 1000 samples, corresponding to 1 s.

To quantify the AIPR elicited by the avatar’s forward-leaning movement, the smoothed COP signal was standardized using the mean and standard deviation of the baseline period from 5.5 to 6 min. This baseline-based standardization was used to account for individual differences in baseline COP position and variability while preserving the direction and relative magnitude of movement-related anterior–posterior displacement in standardized units. Standardization of COP measures has previously been used to facilitate comparisons of postural responses across participants (Pyasik et al., 2026). AIPR was then computed as the mean standardized COP value during the 30-s period after the avatar’s forward-leaning movement. Positive AIPR values indicated anterior COP displacement after the avatar’s forward-leaning movement, whereas negative values indicated posterior displacement.

#### Heart rate

Electrocardiogram (ECG) data were used to extract R–R intervals, from which instantaneous heart rate (IHR) was calculated as beats per minute. IHR was analyzed to assess event-related changes in heart rate associated with the avatar’s hand movement and the avatar’s forward-leaning movement.

Heart-rate variability was also analyzed in the frequency domain. To examine time-varying changes in autonomic activity, time–frequency analysis was applied to the heart rate signal using wavelet transform. Power was estimated over time in the low-frequency (LF: 0.04–0.15 Hz) and high-frequency (HF: 0.15–0.40 Hz) bands, following previous work using time–frequency analysis of heart rate variability to assess dynamic autonomic changes (Tan et al., 2003). The LF/HF ratio was then calculated at each time point and used as an exploratory heart-rate variability index.

To capture event-related changes in IHR and LF/HF ratio, the mean value during the corresponding 10-s pre-event baseline period was subtracted from the mean value during the 10-s post-event period for each measure. A 10-s window was selected to summarize short-latency changes in IHR around each avatar movement. This analysis window was selected given previous psychophysiological evidence showing that heart-rate responses to novel, salient, or unexpected events can emerge within the first several seconds after stimulus onset (Bradley et al., 2012; Noordewier et al., 2021; Zimmer & Richter, 2023). For LF/HF ratio, time-varying LF/HF estimates were first obtained across the entire heart-rate time series, and event-related changes were then calculated using the same 10-s pre- and post-event windows. For the avatar hand-movement event, which occurred 3 min after the start of the experiment, the pre-event baseline period was defined as 170–180 s, and the post-event period was defined as 180–190 s. For the avatar’s forward-leaning event, which occurred 6 min after the start of the experiment, the pre-event baseline period was defined as 350–360 s, and the post-event period was defined as 360–370 s. These post-minus-pre differences were used as outcome measures.

#### Skin conductance response

Skin conductance responses (SCRs) were analyzed using a model-based general linear model (GLM) approach implemented in PsPM to estimate event-related response amplitudes (Bach et al., 2009). PsPM is a MATLAB-based toolbox for model-based analysis of psychophysiological signals and enables estimation of stimulus-evoked electrodermal responses using a predefined canonical response function (CRF).

In this analysis, event onsets at 3 min and 6 min were specified in the GLM. The 3-min onset corresponded to the avatar hand-movement event in the Disruption condition, whereas the 6-min onset corresponded to the avatar forward-leaning event in both conditions. For each onset, predicted SCR time courses were generated by convolving the event onset with the CRF, and these time courses were entered as regressors in the design matrix. The observed SCR signal was modeled as follows:

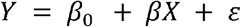

where *Y* represents the observed SCR signal, *pr* represents the intercept, *X* represents the predicted SCR regressor, /? denotes the regression coefficient associated with each experimental event, and ε represents the residual error. Regression coefficients were estimated using the least-squares method. Specifically, /? values were estimated separately for the 3-min and 6-min event onsets. These /? values were used as indices of event-related sudomotor sympathetic activity.

#### Statistical analysis

Questionnaire responses were analyzed using cumulative link mixed models (CLMMs) to examine whether the effect of experimental condition differed between ownership-related and self-location-related aspects of avatar embodiment. CLMMs were adopted because questionnaire responses were measured on a seven-point Likert scale (−3 to +3), and this approach models ordinal outcomes without assuming equal intervals between response categories while accounting for repeated observations within participants through random effects (Hedeker, 2015).

Questionnaire items were initially grouped into ownership-related (Q1–Q3) and self-location-related (Q4–Q5) domains according to their conceptual content. Questionnaire item was included as a fixed effect, and participant was included as a random intercept. To test whether the effect of condition differed between domains, a model assuming a common condition effect across all items was compared with a model estimating separate condition effects for the ownership and self-location domains using a likelihood-ratio test. Condition effects within each domain were reported as odds ratios with 95% confidence intervals, and the p values for the two domain-specific comparisons were adjusted using the Holm method.

Wilcoxon rank-sum tests were used for between-condition comparisons of COP measures, heart-rate-related measures, and SCR indices. One-sample Wilcoxon signed-rank tests were used to examine whether each measure differed from zero within each condition. For each outcome, the p values from the two condition-specific comparisons against zero (Embodiment vs zero and Disruption vs zero) were adjusted using the Holm method. Between-condition comparisons were not adjusted because each involved a single planned comparison within each outcome. Spearman’s rank correlation analyses were performed separately within each condition to examine the association of AIPR with SCR and questionnaire ratings. Because correlations with multiple questionnaire items involved repeated tests of the same subjective measure, p values for AIPR–questionnaire correlations were adjusted within each condition using the Holm method. All statistical analyses were performed using R version 4.5.1. CLMMs were fitted using the clmm function in the ordinal package (version 2025.12-29; Christensen, 2025).

## Results

### Questionnaire

The distributions of questionnaire responses in the Embodiment and Disruption conditions are shown in Fig. 2A. To formally examine whether the effect of experimental condition differed between questionnaire domains, questionnaire responses were analyzed using cumulative link mixed models. The model assuming a common condition effect across all questionnaire items did not show a significant overall effect of condition (likelihood-ratio test: /*^2^*(1) = 1.79, *p* = 0.181). However, allowing the condition effect to differ between the ownership and self-location domains significantly improved model fit *(/^2^* (1) = 5.10, *p* = 0.024).

**Fig. 2.**
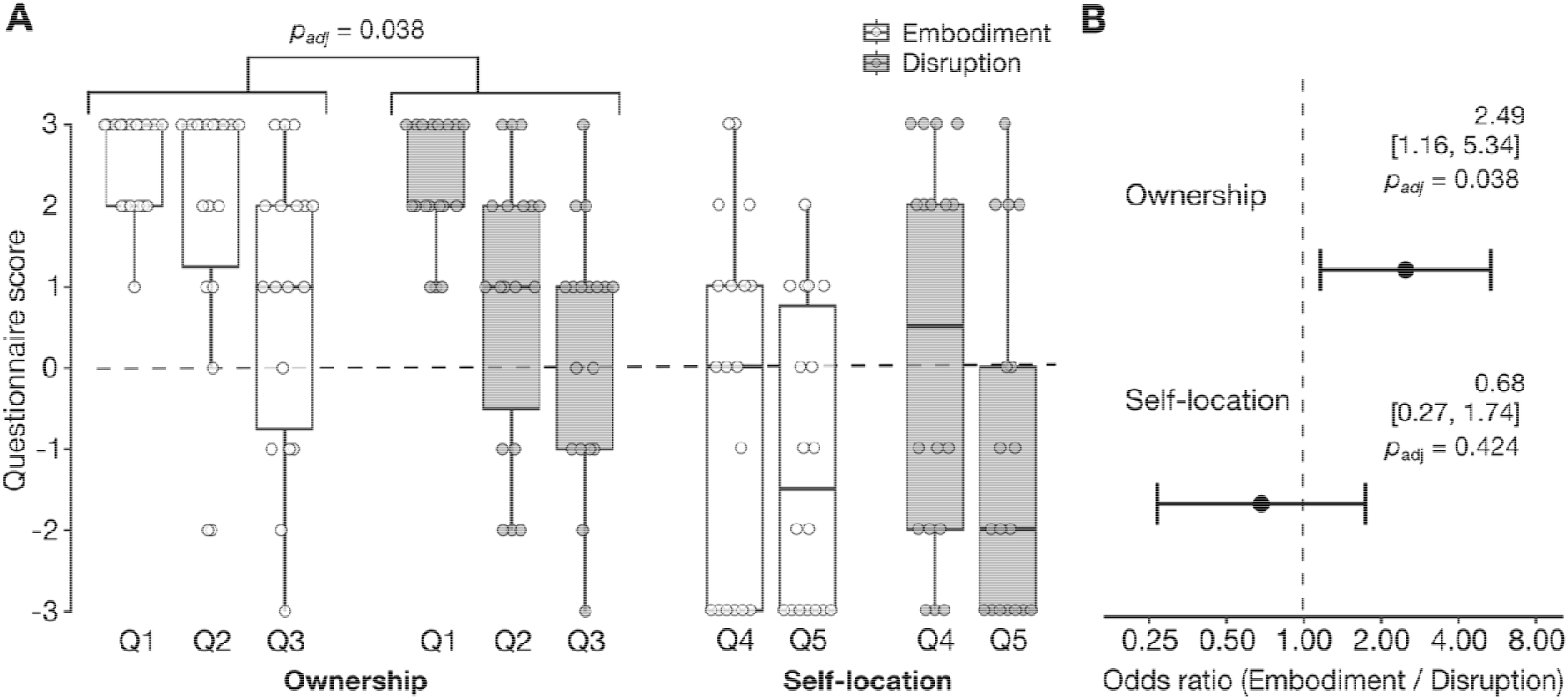
**Questionnaire scores and domain-specific condition effects. A**: Distribution of ownership-related (Q1–Q3) and self-location-related (Q4–Q5) questionnaire scores in the Embodiment and Disruption conditions. Boxes indicate medians and interquartile ranges; whiskers indicate 1.5 × IQR; circles represent individual participants. Brackets indicate the significant domain-level condition effect. **B**: Odds ratios and 95% confidence intervals for the Embodiment condition relative to the Disruption condition estimated from the cumulative link mixed model. The dashed line indicates an odds ratio of 1. *p* values were adjusted using the Holm method.

Ownership-related ratings (Q1–Q3) were significantly higher in the Embodiment condition than in the Disruption condition (odds ratio = 2.49, 95% CI [1.16, 5.34], Holm-adjusted *p* = 0.038), whereas self-location-related ratings (Q4–Q5) did not differ between conditions (odds ratio = 0.68, 95% CI [0.27, 1.74], Holm-adjusted *p* = 0.424). The condition effect was also significantly greater for the ownership domain than for the self-location domain (*z* = 2.24, *p* = 0.025; Fig. 2B).

### COP

Representative COP time courses from the Embodiment and Disruption conditions are shown in Fig. 3A. COP fluctuated continuously throughout the 7-min standing period, reflecting ongoing postural adjustments during standing. The representative traces illustrate the temporal characteristics of COP fluctuations and the pre- and post-event periods used to derive AIPR. To quantify the event-related postural response following the avatar’s forward-leaning movement, the 30-s period immediately preceding the event (330–360 s) was used as the baseline, and the subsequent 30-s period (360–390 s) was defined as the analysis window. The COP signal was standardized relative to the pre-event baseline, and AIPR was calculated as the mean

**Fig. 3.**
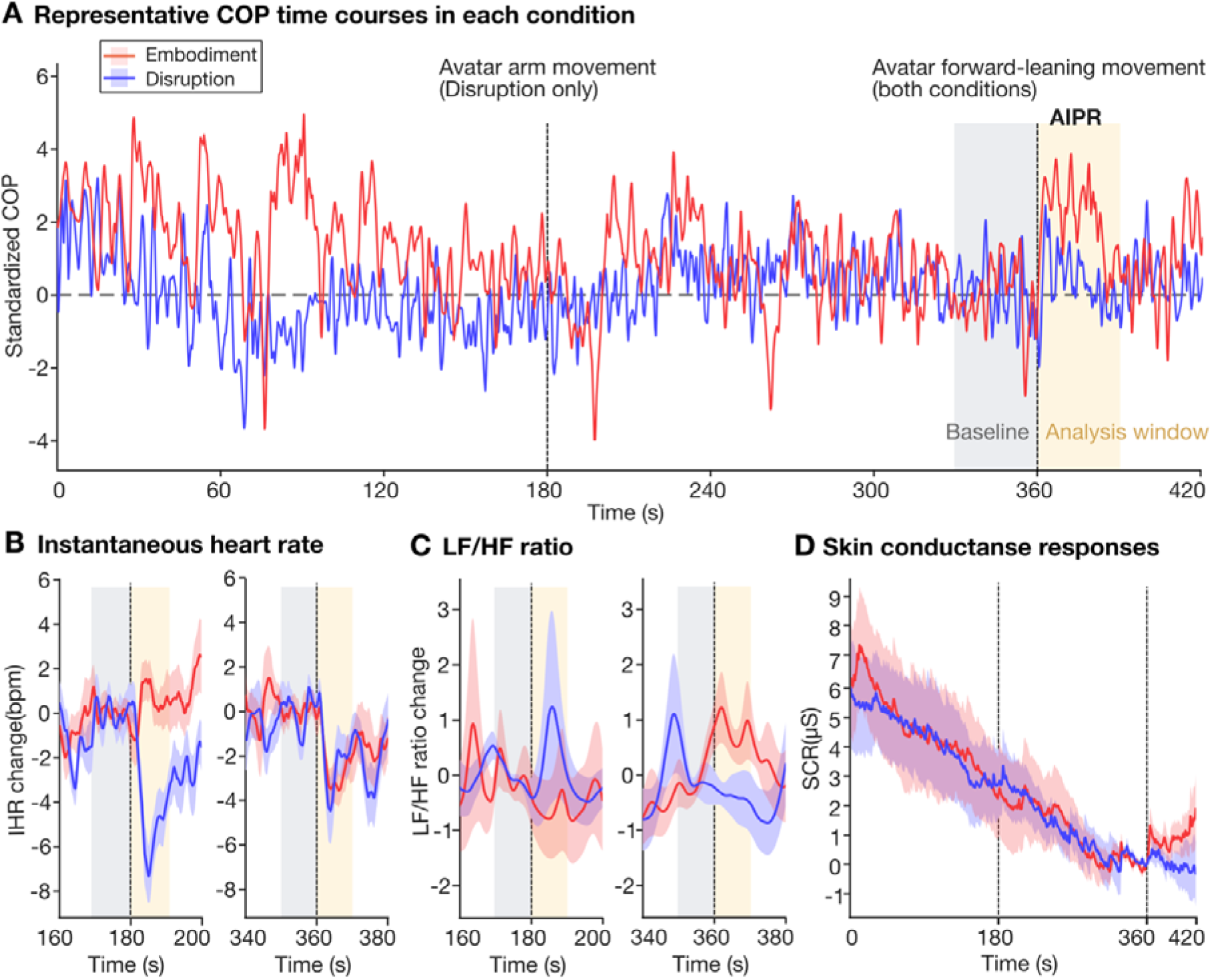
**Time courses of postural and physiological responses during the 7-min VR session. A**: Representative COP time courses from one participant in each condition across the 7-min VR session. The red and blue traces show standardized COP in the Embodiment and Disruption conditions, respectively. The gray shaded region indicates the 30-s baseline period (330–360 s) used for COP standardization, and the beige shaded region indicates the 30-s analysis window (360–390 s) used to calculate AIPR. AIPR was defined as the mean standardized COP value during this post-event analysis window. **B**: Changes in instantaneous heart rate (IHR) around the avatar arm movement (left) and avatar forward-leaning movement (right). **C**: Changes in the LF/HF ratio around the same two events. For panels B and C, vertical dashed lines indicate event onset, gray shaded regions indicate the pre-event baseline periods, and beige shaded regions indicate the post-event analysis periods used in the statistical analyses. **D**: Skin conductance responses (SCRs) across the full experimental session. Vertical dashed lines indicate the event onsets entered into the general linear model at 180 and 360 s. For panels (B–D), solid lines indicate group means and shaded areas indicate ±1 standard error.

#### AIPR

For all one-sample comparisons against zero reported in the following sections, p values were adjusted using the Holm method across the two conditions within each outcome.

AIPR values were significantly greater in the Embodiment condition than in the Disruption condition (*W* = 227.0, *p* = 0.0413, *r* = 0.34; Wilcoxon rank-sum test). One-sample Wilcoxon signed-rank tests showed that AIPR values were significantly greater than zero in the Embodiment condition (*W* = 144, *p* = 0.023, *r* = 0.60), whereas they did not differ significantly from zero in the Disruption condition (*W* = 91.0, *p* = 0.828, *r* = 0.0564). AIPR results are shown in Fig. 4A.

**Fig. 4.**
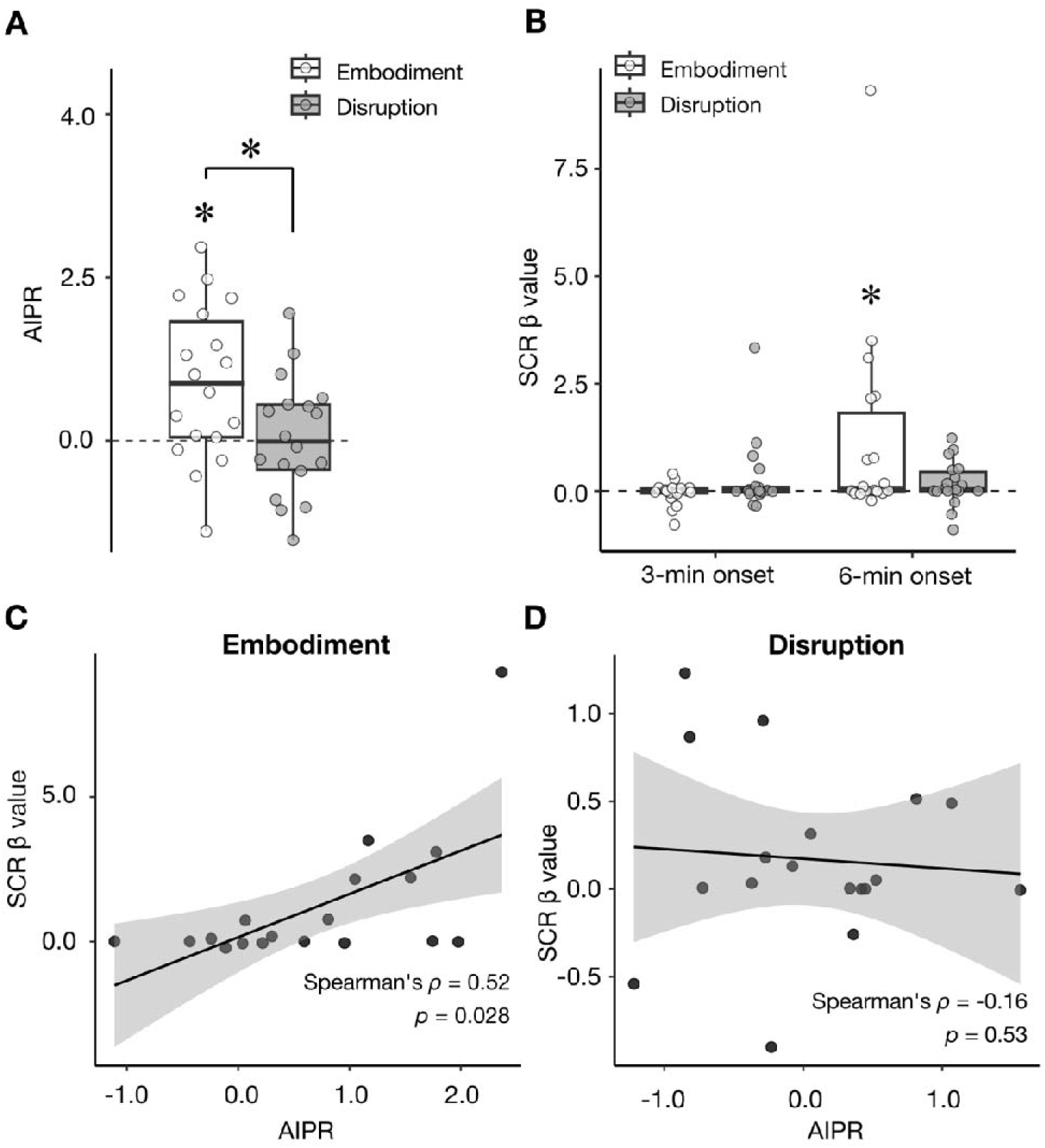
**Postural and autonomic responses associated with avatar movements. A**: AIPR following the avatar’s forward-leaning movement. AIPR was calculated as the mean standardized anterior–posterior COP value during 6.0–6.5 min using the 5.5–6.0-min period as the baseline. **B**: Event-related skin conductance responses, expressed as SCR *fl* values, for the avatar arm movement at 3 min and the avatar forward-leaning movement at 6 min. **C, D**: Associations between AIPR and SCR *fl* values for the forward-leaning movement. Spearman’s rank correlation coefficients were calculated separately for each condition. Regression lines are shown for visualization only. \**p* < 0.05.

### Heart rate

#### Instantaneous heart rate

The time course of changes in instantaneous heart rate is shown in Fig. 3B. During the 180–190 s period following the avatar hand movement, the change in IHR was significantly lower in the Disruption condition than in the Embodiment condition (*W* = 258, *p* = 0.0025, *r* = 0.51; Wilcoxon rank-sum test). One-sample Wilcoxon signed-rank tests showed that the IHR change was significantly lower than zero in the Disruption condition (*W* = 2.00, *p* < 0.001, *r* = 0.85), whereas it did not differ significantly from zero in the Embodiment condition (*W* = 91.0, *p* = 0.828, *r* = 0.056).

During the 360–370 s period following the avatar forward-leaning movement, the IHR change did not differ significantly between the Embodiment and Disruption conditions (*W* = 146, *p* = 0.624, *r* = 0.084; Wilcoxon rank-sum test). However, one-sample Wilcoxon signed-rank tests showed that the IHR change was significantly lower than zero in both the Embodiment condition (*W* = 7.00, *p* = 0.001, *r* = 0.80) and the Disruption condition (*W* = 32.0, *p* = 0.021, *r* = 0.55).

#### LF/HF ratio

The time course of changes in the LF/HF ratio is shown in Fig. 3C. During the 180–190 s period following the avatar hand movement, the LF/HF ratio change did not differ significantly between the Embodiment and Disruption conditions (*W* = 153, *p* = 0.788, *r* = 0.0475; Wilcoxon rank-sum test).

During the 360–370 s period following the avatar forward-leaning movement, the LF/HF ratio change was significantly higher in the Embodiment condition than in the Disruption condition (*W* = 259, *p* = 0.002, *r* = 0.51; Wilcoxon rank-sum test). One-sample Wilcoxon signed-rank tests showed that the LF/HF ratio change was significantly greater than zero in the Embodiment condition (*W* = 151, *p* = 0.009, *r* = 0.672), whereas it did not differ significantly from zero in the Disruption condition (*W* = 48.0, *p* = 0.107, *r* = 0.385).

### Skin conductance response

The time course of skin conductance responses (SCRs) is shown in Fig. 3D. β values estimated for the 3-min and 6-min event onsets did not differ significantly between the Embodiment and Disruption conditions (3-min onset: *W* = 128, *p* = 0.289, *r* = 0.179; 6-min onset: *W* = 188, *p* = 0.420, *r* = 0.137; Wilcoxon rank-sum tests). In within-condition analyses, β values estimated for the onset of the avatar forward-leaning event did not differ significantly from zero in either the Embodiment condition (*W* = 135, *p* = 0.066, *r* = 0.508) or the Disruption condition (*W* = 124, *p* = 0.0979, *r* = 0.395). These β-value results are shown in Fig. 4B.

### Correlation analyses

To examine the relationship between AIPR and SCR responses to the avatar’s forward-leaning movement, Spearman’s rank correlation coefficients were calculated separately for each condition. A significant positive correlation was observed in the Embodiment condition *(ρ* = 0.52, *p* = 0.028, Fig.4C), whereas no significant correlation was found in the Disruption condition *(ρ* = −0.16, *p* = 0.53, Fig.4D).

No significant correlations were found between AIPR and questionnaire scores after Holm correction (Table S1).

## Discussion

The present study examined whether avatar-related body ownership modulates participants’ postural responses to an unexpected change in avatar posture. AIPR was used as an event-related behavioral measure of anterior–posterior COP displacement elicited by a forward-leaning movement of a virtual avatar. The main finding was that AIPR values were significantly greater in the Embodiment condition than in the Disruption condition. Moreover, AIPR values were significantly greater than zero only in the Embodiment condition, suggesting that participants showed a forward postural response when ownership-related body–avatar correspondence was maintained. AIPR values were also positively correlated with SCR β values in the Embodiment condition. In contrast, AIPR values did not show a significant correlation with subjective questionnaire scores. Together, these findings suggest that avatar-related embodiment can be expressed not only in subjective reports, but also in the participant’s postural regulation, which was further associated with autonomic responses.

The questionnaire results indicate that the experimental manipulation selectively influenced ownership-related aspects of avatar embodiment. Although no significant overall condition effect was observed across questionnaire items, the cumulative link mixed model revealed that the effect of condition differed between questionnaire domains. Specifically, ownership-related ratings (Q1–Q3) were significantly higher in the Embodiment condition than in the Disruption condition, whereas self-location-related ratings (Q4–Q5) did not differ between conditions. These findings suggest that the Disruption manipulation primarily affected the sense of ownership over the avatar rather than the perceived location of the self. This interpretation is consistent with previous full-body illusion studies showing that body ownership is more susceptible than self-location to manipulations that disrupt body–avatar correspondence. For example, body ownership is reduced when the avatar’s body is tilted relative to the participant’s orientation and gravity (Thür et al., 2019). Taken together, the questionnaire findings indicate that the Embodiment condition maintained stronger ownership-related body–avatar correspondence than the Disruption condition.

The central implication of the AIPR finding is that an avatar’s posture can influence the participant’s own postural control when ownership-related body–avatar correspondence is maintained. Pyasik et al. (2026) showed that when a virtual body was embodied from the first-person perspective, participants’ COP shifted in accordance with the flexion and extension phases of the avatar’s repeated squats, whereas this phase-locked postural modulation was absent in the third-person perspective condition. Subjective embodiment was also present only in the first-person perspective condition, leading the authors to interpret the absence of COP modulation in the third-person perspective condition as supporting a role of embodiment rather than action observation alone. This suggests that postural control can be modulated by avatar movement when the virtual body is experienced as one’s own. A related effect has also been reported in the rubber hand illusion: movements of an embodied rubber hand elicited corresponding spontaneous movements of the participant’s own hand (Shibuya et al., 2018). These findings suggest that changes occurring in an embodied artificial body can be reflected back onto the participant’s own body, a phenomenon that has been discussed as *back-projection* of bodily self-representation (Shimada, 2022; Shimada et al., 2026). In the present study, participants viewed the back of an avatar rather than seeing the avatar as a direct visual substitute for their own body. Nevertheless, synchronous visuo-tactile stimulation and body-size matching were used to support body–avatar correspondence, and the Disruption condition transiently violated this correspondence before the avatar’s forward-leaning movement. The larger AIPR values in the Embodiment condition suggest that the avatar’s forward-leaning movement influenced the participant’s postural control when ownership-related bodily correspondence was maintained. Taken together, these findings indicate that avatar-induced postural responses depend not merely on visual observation of avatar movement, but on whether body–avatar correspondence allows the avatar to be incorporated into the participant’s bodily representation.

AIPR values did not show a significant correlation with questionnaire scores, even though the experimental manipulation significantly increased ownership-related questionnaire ratings at the group level. One possible interpretation is that AIPR captures an aspect of avatar-related embodiment expressed through postural control that is not fully reflected in explicit questionnaire items. This interpretation is consistent with previous bodily illusion research showing that different indices of embodiment do not necessarily covary. In the rubber hand illusion, proprioceptive drift, defined as a shift in the perceived position of the participant’s own hand toward the rubber hand, has been widely used as a behavioral measure of illusion-related bodily recalibration (Botvinick & Cohen, 1998). However, subsequent studies have shown that subjective ownership ratings and proprioceptive drift do not always correspond, suggesting that they reflect partly distinct aspects of bodily illusion (Rohde et al., 2011; Shimada et al., 2014). Similarly, full-body illusion studies have shown that body ownership and self-location can be dissociated (Maselli & Slater, 2014). Thus, the absence of a significant correlation between AIPR values and questionnaire scores does not necessarily indicate that AIPR is unrelated to avatar-related embodiment. Rather, it may suggest that AIPR captures a different, postural-control aspect of embodiment.

In this respect, AIPR may be conceptually analogous to proprioceptive drift in the rubber hand illusion, while differing in the bodily process it captures. Proprioceptive drift reflects a recalibration of perceived limb position, whereas AIPR reflects an event-related change in postural regulation induced by the movement of an embodied avatar. Although AIPR involves anterior–posterior displacement, it differs from conventional self-location measures in that it quantifies an actual COP-based postural adjustment rather than the perceived location of the self. Therefore, AIPR should not be interpreted as a direct objective measure of subjective full-body illusion strength. Rather, it may provide an objective behavioral measure of the extent to which an avatar-related bodily representation influences the participant’s own postural control. Accordingly, AIPR should be interpreted alongside subjective ratings, behavioral measures of self-location, and autonomic responses, rather than as a replacement for them.

The association between AIPR and SCR further supports the idea that AIPR reflects more than a nonspecific fluctuation in standing posture. Previous full-body illusion studies have shown that threatening or aversive events directed toward an embodied body can elicit autonomic responses such as increased SCR (Ehrsson, 2007; Petkova & Ehrsson, 2008). In the present study, AIPR values were positively correlated with SCR β values in the Embodiment condition. Thus, participants who showed stronger forward postural displacement also tended to show stronger autonomic responses to the avatar’s postural change. Although SCR did not significantly differ between the Embodiment and Disruption conditions, the positive association between AIPR and SCR observed in the Embodiment condition suggests that the postural response was associated with autonomic activity when ownership-related body–avatar correspondence was maintained. This pattern supports the interpretation of AIPR as a complementary behavioral measure of avatar-related embodiment, while also indicating that it should be interpreted together with subjective and physiological indices rather than as a standalone measure.

Heart-rate-related measures also showed event-related changes, although these findings should be considered exploratory. IHR decreased in the Disruption condition after the avatar’s hand movement and in both conditions after the avatar’s forward-leaning movement. Previous full-body illusion studies have reported heart-rate deceleration in response to threatening or aversive changes directed toward an embodied virtual body (Slater et al., 2010; Maselli & Slater, 2013). The present IHR decreases may therefore reflect a broader orienting or aversive response to unexpected changes in the avatar. In addition, the LF/HF ratio change following the avatar’s forward-leaning movement was greater in the Embodiment condition than in the Disruption condition and was significantly greater than zero only in the Embodiment condition. This condition-dependent modulation suggests that cardiovascular responses to the avatar’s postural change may also be influenced by body–avatar correspondence.

However, given the exploratory nature of these analyses and the different response patterns observed across the heart-rate-related measures, the functional significance of these cardiovascular changes remains unclear. Future studies with hypotheses specifically targeting cardiovascular responses are needed to determine how IHR and LF/HF changes relate to avatar embodiment.

Beyond its theoretical implications, the methodological significance of AIPR lies in its ability to capture an event-related postural component of avatar-related embodiment while participants remain standing in place. Existing behavioral measures of bodily illusion, such as proprioceptive drift in the rubber hand illusion or self-location drift in full-body illusion, primarily assess perceived body or self-location. In contrast, AIPR is derived from continuous COP recordings and therefore indexes how the participant’s postural control responds to a sudden change in an avatar that is subjectively linked to their own body. Because it does not require locomotion or explicit spatial judgments during the illusion, AIPR may provide a useful event-related measure of bodily responses in VR without interrupting the ongoing illusion.

This may be particularly relevant for situations in which the avatar is linked to the user’s body through visuotactile correspondence or other ownership-related body–avatar mappings. For example, VR systems designed to guide posture or movement may benefit from measures that capture how strongly avatar movements influence the user’s own bodily control. The present study provides an initial step by showing that AIPR values differed between Embodiment and Disruption conditions and were positively associated with SCR β values in the Embodiment condition. Future studies should examine the reliability, specificity, and generalizability of AIPR across tasks, populations, avatar configurations, and avatar movements. It would also be important to determine whether higher-level contextual factors can modulate avatar-induced postural responses. Recent work has shown that narrative information about an avatar can influence the sense of agency toward that avatar (Hamagashira et al., 2026), while conceptual accounts of narrative embodiment propose that higher-level self-related information may ultimately shape embodied behavior (Shimada et al., 2026). Future studies could examine whether higher-level contextual information about an avatar, such as narrative framing, can influence avatar-induced postural responses in addition to sensorimotor body–avatar correspondence.

Several limitations should be noted. First, the present study used a between-subject design. Although participants were randomly assigned to the Embodiment and Disruption conditions, individual differences in postural control, susceptibility to body illusions, and autonomic reactivity may have influenced the results. Future studies using a within-subject design would allow stronger conclusions about how changes in avatar embodiment modulate AIPR values within the same individuals. Second, all participants were male because the avatar used in the present experiment was male. This limits the generalizability of the findings. Future studies should examine whether similar postural responses are observed when the participant’s gender and the avatar’s gender are systematically manipulated. Third, the Disruption manipulation may have affected not only body ownership but also attention, surprise, prediction error, or the perceived naturalness of the avatar’s movement. Therefore, the difference in AIPR values between conditions cannot be attributed exclusively to body ownership. More refined control conditions will be needed to dissociate the effects of ownership-related body–avatar correspondence from the effects of unexpected avatar movement. Fourth, although AIPR values were associated with SCR in the Embodiment condition, the sample size was relatively small for correlation analyses. Therefore, the relationship between postural and autonomic responses should be replicated in larger samples. Future studies should also examine whether AIPR is observed for other avatar movements and whether the direction and magnitude of COP displacement systematically correspond to the avatar’s postural changes. Finally, the present study did not clarify the neural or motor-control mechanisms by which avatar posture influences the participant’s postural control. Future studies combining AIPR assessment with neurophysiological measures may help clarify how avatar-related embodiment affects postural regulation.

## Conclusion

The present study proposes AIPR as a COP-based event-related measure for examining postural responses to avatar movement in the context of avatar-related embodiment. The findings suggest that an avatar’s postural change can influence the participant’s own postural regulation when ownership-related body–avatar correspondence is maintained. Analogous to proprioceptive drift in the rubber hand illusion, AIPR may provide an objective behavioral index of a bodily response associated with avatar-related embodiment. However, AIPR does not directly measure subjective full-body illusion strength. Rather, it may capture a postural-control component of avatar embodiment that is partly dissociable from explicit subjective reports. By complementing subjective ownership ratings, autonomic responses, and self-location measures, AIPR may provide a practical tool for investigating how embodied avatars influence the participant’s bodily state in virtual reality.

## Acknowledgements

Generative AI was used solely for English-language editing and proofreading. The authors used Codex (GPT-5.6, OpenAI, https://openai.com). All AI-assisted suggestions were reviewed and revised by the authors, who take full responsibility for the accuracy and integrity of the manuscript.

## Author contributions

YT: Formal analysis, Visualization, Writing – original draft; KF: Formal analysis, Investigation, Methodology; KH: Supervision, Writing – review & editing; SS: Conceptualization, Supervision, Writing – original draft, Writing – review & editing.

## Data availability

The data that support the findings of this study are available from the corresponding author upon reasonable request.

