## Supplementary material for "Avatar-Induced Postural Responses during a Virtual Full-Body Illusion": Table S1

**Supplementary correlation analyses**

Spearman’s rank correlation analyses were conducted separately for the Embodiment and Disruption conditions. For associations between AICD and questionnaire items (Q1–Q5), p values were adjusted within each condition using the Holm method. No significant correlations were observed between AICD and any questionnaire item after correction for multiple comparisons (Table S1).

**Table S1. Spearman correlations between AICD and questionnaire items.**

|  | ***ρ*** | ***p*** | ***p*_adj._** |
| --- | --- | --- | --- |
| ***Embodiment*** |  |  |  |
| Q1 | 0.061 | 0.809 | 1.000 |
| Q2 | 0.276 | 0.267 | 1.000 |
| Q3 | 0.019 | 0.941 | 1.000 |
| Q4 | -0.282 | 0.257 | 1.000 |
| Q5 | 0.130 | 0.608 | 1.000 |
| ***Disruption*** |  |  |  |
| Q1 | 0.145 | 0.565 | 1.000 |
| Q2 | -0.195 | 0.438 | 1.000 |
| Q3 | 0.119 | 0.639 | 1.000 |
| Q4 | -0.347 | 0.158 | 1.000 |
| Q5 | -0.239 | 0.339 | 1.000 |
